# Calibration of FRET-based biosensors using multiplexed biosensor barcoding

**DOI:** 10.1101/2024.09.04.610346

**Authors:** Jhen-Wei Wu, Jr-Ming Yang, Chao-Cheng Chen, Gabriel Au, Suyang Wang, Gia-Wei Chern, Chuan-Hsiang Huang

## Abstract

Förster resonance energy transfer (FRET) between fluorescent proteins (FPs) is widely used in the design of genetically encoded fluorescent biosensors, which are powerful tools for monitoring the dynamics of biochemical activities in live cells. FRET ratio, defined as the ratio between acceptor and donor signals, is often used as a proxy for the actual FRET efficiency, which must be corrected for signal crosstalk using donor-only and acceptor-only samples. However, the FRET ratio is highly sensitive to imaging conditions, making direct comparisons across different experiments and over time challenging. Inspired by a method for multiplexed biosensor imaging using barcoded cells, we reasoned that calibration standards with fixed FRET efficiency can be introduced into a subset of cells for normalization of biosensor signals. Our theoretical analysis indicated that the FRET ratio of high-FRET species relative to non-FRET species slightly decreases at high excitation intensity, suggesting the need for calibration using both high and low FRET standards. To test these predictions, we created FRET donor-acceptor pairs locked in “FRET-ON” and “FRET-OFF” conformations and introduced them into a subset of barcoded cells. Our results confirmed the theoretical predictions and showed that the calibrated FRET ratio is independent of imaging settings. We also provided a strategy for calculating the FRET efficiency. Together, our study presents a simple strategy for calibrated and highly multiplexed imaging of FRET biosensors, facilitating reliable comparisons across experiments and supporting long-term imaging applications.

## Background

Cellular functions depend on precisely coordinated activities of large numbers of biomolecules in space and time. To understand physiological and pathological processes in cells, it is often necessary to track the spatiotemporal dynamics of molecular activities as well as their responses to perturbations. Genetically encoded fluorescent biosensors, which incorporate fluorescent proteins (FPs) that alter their emission properties or subcellular localization in response to specific molecular activations or changes in the cell’s physicochemical environment, are powerful tools for monitoring these activities [1–3].

Unlike methods that require cell lysis or fixation, such as mass spectrometry or antibody-based techniques, biosensors offer several key advantages. For example, they enable continuous monitoring of molecular activities in live cells and can be targeted to specific subcellular compartments or organelles, providing localized reports of activity. Additionally, biosensors reveal cell-to-cell variability that might be masked in ensemble measurements. Moreover, some biochemical events such as the activation of G-proteins involve subtle conformational changes that are hard to detect with other approaches but can be readily monitored using biosensors [4]. Beyond molecular activities, some biosensors can monitor the physical environment of the cell, including pH and membrane potential [5,6].

Förster Resonance Energy Transfer (FRET) is a process involving non-radiative energy transfer from a donor fluorophore to a nearby acceptor fluorophore and is extensively used in the design of biosensors [7,8]. In FRET-based biosensors, binding of a target molecule to the recognition element triggers conformational changes, altering the orientation or distance between the donor and acceptor fluorophores. This leads to a change in FRET efficiency, which serves as a readout for the target activity. For example, many kinase biosensors use a substrate specific to the kinase of interest, where phosphorylation of the substrate causes a conformational shift that affects the FRET efficiency between two FPs. The change in FRET efficiency provides a quantitative measure of kinase activity [9].

There are several methods to measure FRET efficiency, including spectral imaging, acceptor photobleaching, and donor fluorescence lifetime imaging [10]. The most commonly used method, sometimes referred to as sensitized emission FRET, involves measuring the emission of both the donor and acceptor under excitation. Ideally, the signal in the acceptor channel should be entirely due to FRET. However, the acceptor can also be directly excited by the laser, albeit at a lower intensity, and donor fluorescence can bleed through into the acceptor channel. To correct these issues, parameters for signal crosstalk must be determined using donor- and acceptor-only samples imaged at different wavelengths [11,12]. This process is not only time-consuming but also prone to errors across multiple measurements. As a result, the acceptor-to-donor signal ratio, or FRET ratio, is often used as a convenient but imperfect surrogate for FRET efficiency. However, this ratio is influenced by imaging conditions, such as laser intensity and detector sensitivity settings, making it difficult to directly compare results across different imaging sessions. Consequently, most FRET biosensor experiments are confined to a single imaging session of limited duration, typically lasting up to several hours.

Most FRET biosensors use CFP and YFP or their variants as the FRET donor and acceptor, respectively, although other FRET pairs such as GFP-RFP have gained increasing popularity in recent years [13]. Due to the similar emission spectrum, fluorescent biosensors are hard to multiplex [14–16]. We recently developed a multiplexed biosensor barcoding method that enables simultaneous imaging of large numbers of biosensors. In this method, cells expressing different biosensors are labeled with a pair of barcoding proteins, which are blue or red fluorescent proteins (FPs) targeted to different subcellular locations. The spectra of barcoding proteins can be separated from those of commonly used biosensors based on GFP, CFP, and YFP. Barcoded cells expressing different biosensors are mixed for simultaneous imaging, and the identity of each biosensor in each cell is established by analyzing the barcode through machine learning models [17–19].

Using the biosensor barcoding method, we reasoned that calibration standards can be expressed in a subset of barcoded cells for normalizing the FRET ratio of biosensors to account for imaging variability. Our theoretical analysis indicated that the normalized FRET ratio slightly decreases with higher excitation intensity, but this can be corrected by calibrating against standards with both low and high FRET efficiency. To test this approach, we created calibration standards using the CFP-YFP FRET pairs locked in “FRET-ON” and “FRET-OFF” conformations. Our results confirmed the theoretical predictions and showed that the calibrated FRET ratio is independent of imaging settings. Moreover, simultaneous imaging of CFP and YFP allows for the calculation of actual FRET efficiency. Together, our study presents a simple and effective strategy for calibrated and multiplexed imaging of FRET biosensors, facilitating reliable comparisons across experiments and supporting long-term imaging applications.

## Results

### Theoretical modeling of FRET calibration

Here we focus on unimolecular FRET biosensors consisting of a single donor and acceptor pair, which is the most commonly used design. We first outline the kinetic theory of FRET [20,21] for the simple case of a single donor and a single acceptor molecule. We assume a two-level system to describe both the donor (D) and acceptor (A) molecules. We use P_X_ and P_X_∗ to denote the probability that molecule species X (here X = D or A) is in the ground or excited state, respectively. Probability conservation of the two-level system implies that P_X_ + P_X_∗ = 1. The time evolution of these probabilities is described by a set of coupled rate equations:

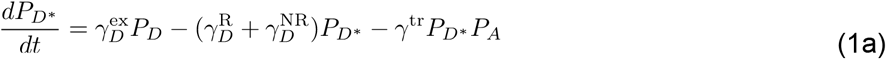

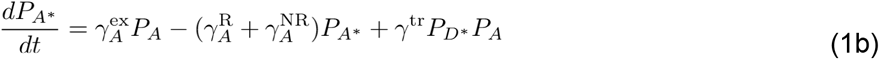

Here 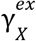 is the rate of excitation of fluorescent species X initially in their ground state, 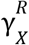 and 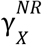 denote the rates of excited-state depopulation through nonradiative (e.g., internal conversion) and radiative (i.e., emission of a photon) processes, respectively. The FRET rate from donor to acceptor is represented by γ^tr^, which is dependent on the orientation and separation of the two fluorescent species. The net energy transfer is also proportional to the probability of the donor being in the excited states as well as the probability that the acceptor is in the ground state. The various electron transitions are summarized in **Fig. 1A**. Numerical simulations of the probabilities of the donor and acceptor in the excited state over time is shown in **Fig. 1B**.

**Figure 1.**
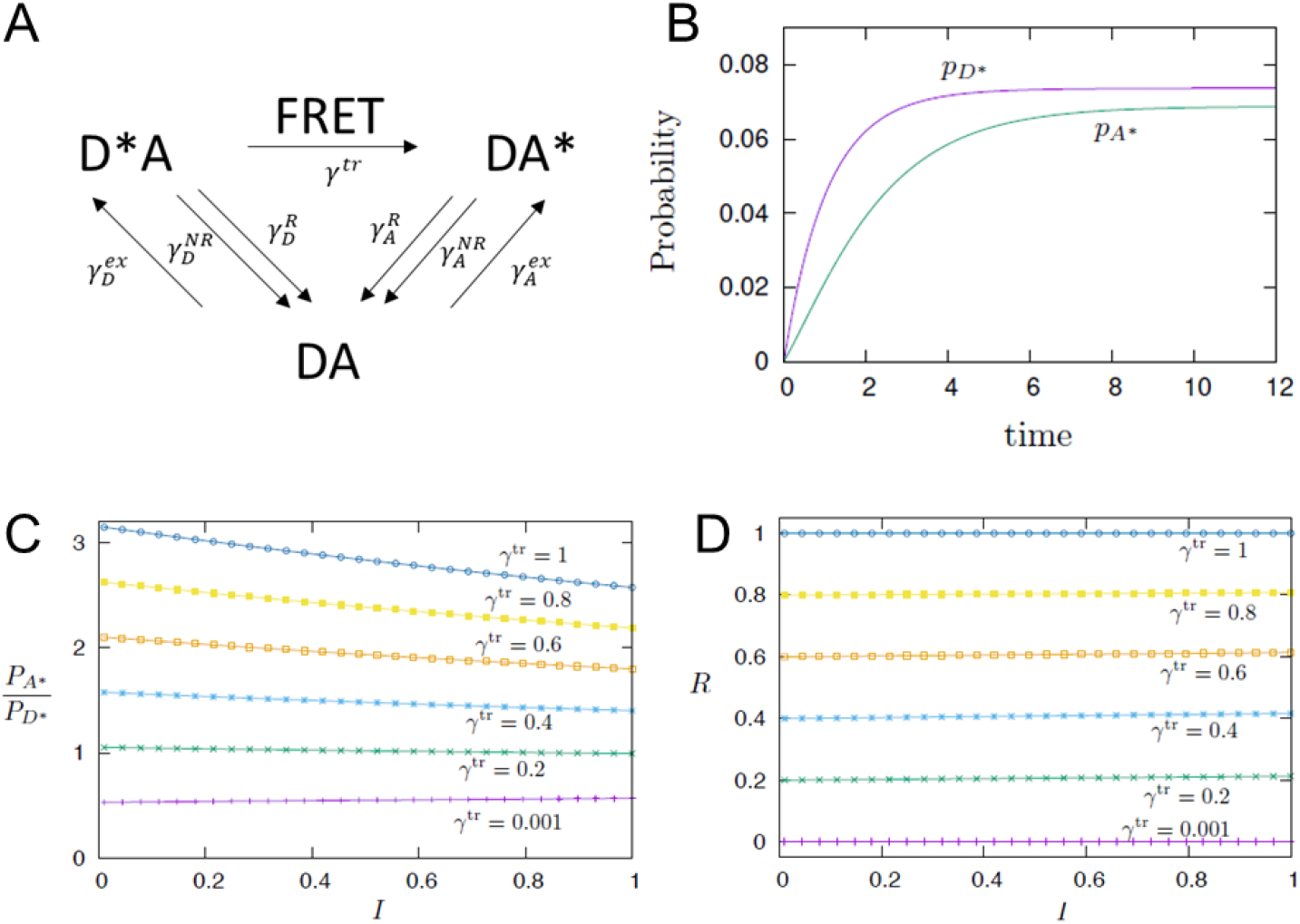
Theoretical modeling. (A) Schematic diagram of the various transitions in a FRET system with one donor and one acceptor molecule. D and A denote donor and acceptor molecules, respectively. Asterisk denotes excited species. Simulated FRET calibration. (B) Probabilities of the donor and acceptor in the excited state over time. The parameters used in the simulations are 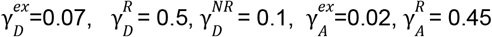, 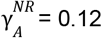 and 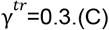 Ratio of the probability of finding donor in the excited state to that of acceptor in the steady state versus laser intensity I at steady state for varying FRET rate γ^tr^. (D) Calibrated ratio of the same simulation data based on γ_1_^*tr*^ =0.001 and γ_2_^*tr*^ = 1.

For single-photon excitation, the excitation rates of each fluorescent species is given by

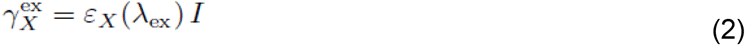

where *I* is the laser irradiance, and ε_X_(λ_ex_) is the wavelength-dependent extinction coefficient, with λ_ex_ being the excitation wavelength.

For biosensor applications, continuous wave (CW) light sources are often employed, indicating a constant laser intensity *I*. We are interested in steady-state solutions, which can be obtained by setting dP_D_∗/dt = dP_A_∗/dt = 0 in Eq. (1). However, in the presence of FRET, the resultant coupled nonlinear equations cannot be solved analytically. Numerical integration of the differential equations is used to obtain the steady state solutions.

The fluorescent signal *S*_*X*_is proportional to the number of photons 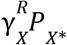 emitted per unit time through the radiative transition, as well as the number *N* of fluorescent molecules:

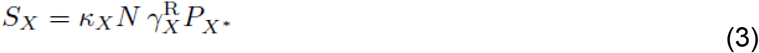

where the species-dependent coefficient *K*_*X*_is determined by the settings of image acquisition. For FRET-based biosensors, we are interested in the ratio of the acceptor to donor fluorescent signals

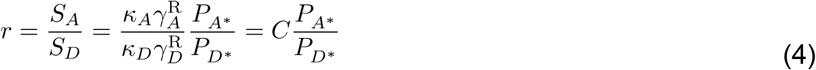

Here the proportional coefficient *C* depends on the excitation and emission wavelengths and settings of image acquisition, but is independent of the laser intensity *I* or the FRET transfer rate γ^tr^. On the other hand, the ratio of the excited state probability depends nontrivially on both *I* and γ^tr^.

**Fig. 1C** shows the probability ratio P_A_∗/P_D_∗ at the steady state as a function of laser intensity *I* for varying FRET transfer rate γ^tr^. The parameters used in the simulations are 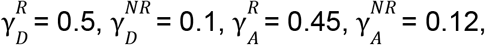, and extinction coefficients ε_D_ = 0.1 and ε_A_ = 0.05. With a small γ, P_A_∗/P_D_∗ stayed relatively unchanged with increasing laser intensity. However, a more pronounced dependence on laser intensity is obtained for a larger FRET rate.

Here we present a calibration method to minimize the dependence of the ratio of acceptor to donor signal on the laser intensity I, while also removing other extrinsic factors related to image acquisition. We select two reference FRET rates, γ_1_ ^tr^ and γ_2_ ^tr^, and define a normalized ratio as

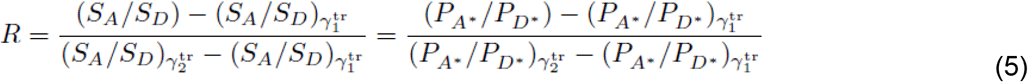

where the subscript 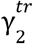 and 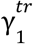 indicates quantities obtained from the two reference FRET coefficients. The calibrated ratios based on the same simulation data are shown in **Fig. 1D**. The results show a negligible dependence on the laser intensity for all FRET rates.

### Linear approximation

To gain further insight into the basis of the calibration formula, we considered linear approximation to the steady state solutions of the FRET equation. As demonstrated in Ref. [21], the rate-equation modeling based on linear approximation agrees well with the experimental results for a CFP-YFP system. Assuming a small excitation probability for the acceptor P_A_∗ ≪ 1, the nonlinear term in Eq. (1) can be approximated as P_D_∗P_A_ = P_D_∗(1 − P_A_∗) ≈ P_D_∗. We first obtain the excitation probability for the donor

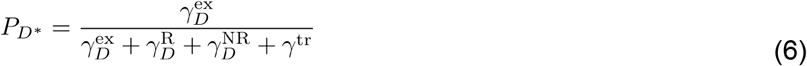

We consider a simplified situation of weak excitation, i.e. γ_D_ ^ex^ ≪ γ_D_ ^R^, γ_D_ ^NR^, and neglect the excitation rate in the denominator. In the absence of FRET, the number of photons emitted per unit time is given by

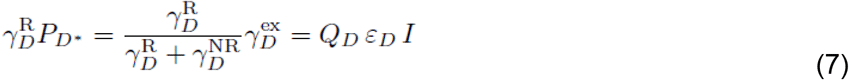

where Φ_D_ = γ_D_ ^R^/(γ_D_ ^R^ + γ_D_ ^NR^) is the quantum yield of the donor molecule. The fluorescent signal is thus proportional to the product of the quantum yield and the extinction ratio. Similar expression also applies to the acceptor molecule. The steady-state solution for P_A_∗ in the linear approximation is

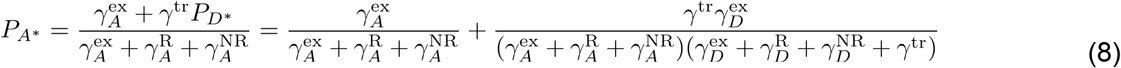

The ratio of the two excitation probabilities is

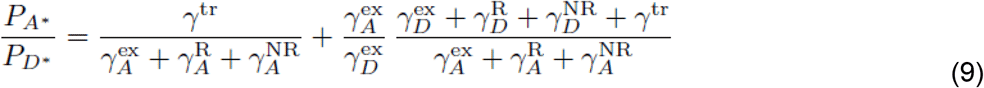

Note that 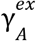 and 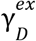 are both proportional to the laser intensity *I*, while 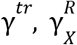 and 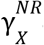 are independent of *I*.

The first term explains the inverse relationship between P_*A*_∗/P_D_∗ and *I* at high levels of γ^tr^ (**Fig. 1C**).

Substitute (9) into the calibrated ratio, we obtain

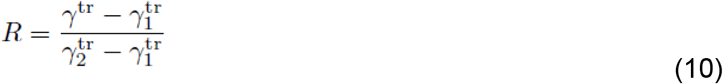

Importantly, this calibrated ratio in the limit of linear approximation is now independent of laser intensity or even the parameters that are related to image acquisition.

### Generation of FRET-ON and FRET-OFF calibration standards

Our theoretical analysis indicates that normalizing the FRET ratio using low and high FRET calibration standards results in a value that depends solely on the FRET efficiency, independent of the excitation intensity. To test this prediction, we generated calibration standards with low and high FRET efficiency. A common biosensor design utilizes CFP (cyan fluorescent protein) as the donor and YFP (yellow fluorescent protein) as the acceptor. Various CFP and YFP variants are available with similar excitation and emission spectra but differing in brightness. Therefore, we generated two calibration constructs that lock the CFP-YFP pairs in the FRET (FREF-ON) or non-FRET (FRET-OFF) configuration.

To generate the FRET-ON construct, we started with a biosensor that undergoes loss of FRET upon target activation. One such biosensor is the CytoFAK biosensor, which contains a FAK substrate sequence [22] (**Fig. 2A**). Phosphorylation of the substrate on the tyrosine residue by activated FAK induces its binding to the SH2 domain, resulting in a conformational change and a corresponding loss of FRET. We mutated the tyrosine in the FAK substrate to phenylalanine (Y349F) to prevent its phosphorylation by FAK (**Fig. 2A**). Interestingly, we noticed that cells expressing CytoFAK showed a lower YFP/CFP in the cytosol than in the nucleus, suggesting partial loss of FRET in the cytosol (**Fig. S1A**). Consistent with this idea, activation and inhibition of FAK caused a decrease and increase, respectively, of YFP/CFP preferentially in the cytosol (**Fig. S1C, D**). In contrast, in cells expressing the FRET-ON construct, YFP/CFP was uniformly high across the cell (**Fig. S1B**) and did not respond to EGF stimulation (**Fig. 2B, C**, and **S1B**).

**Figure 2.**
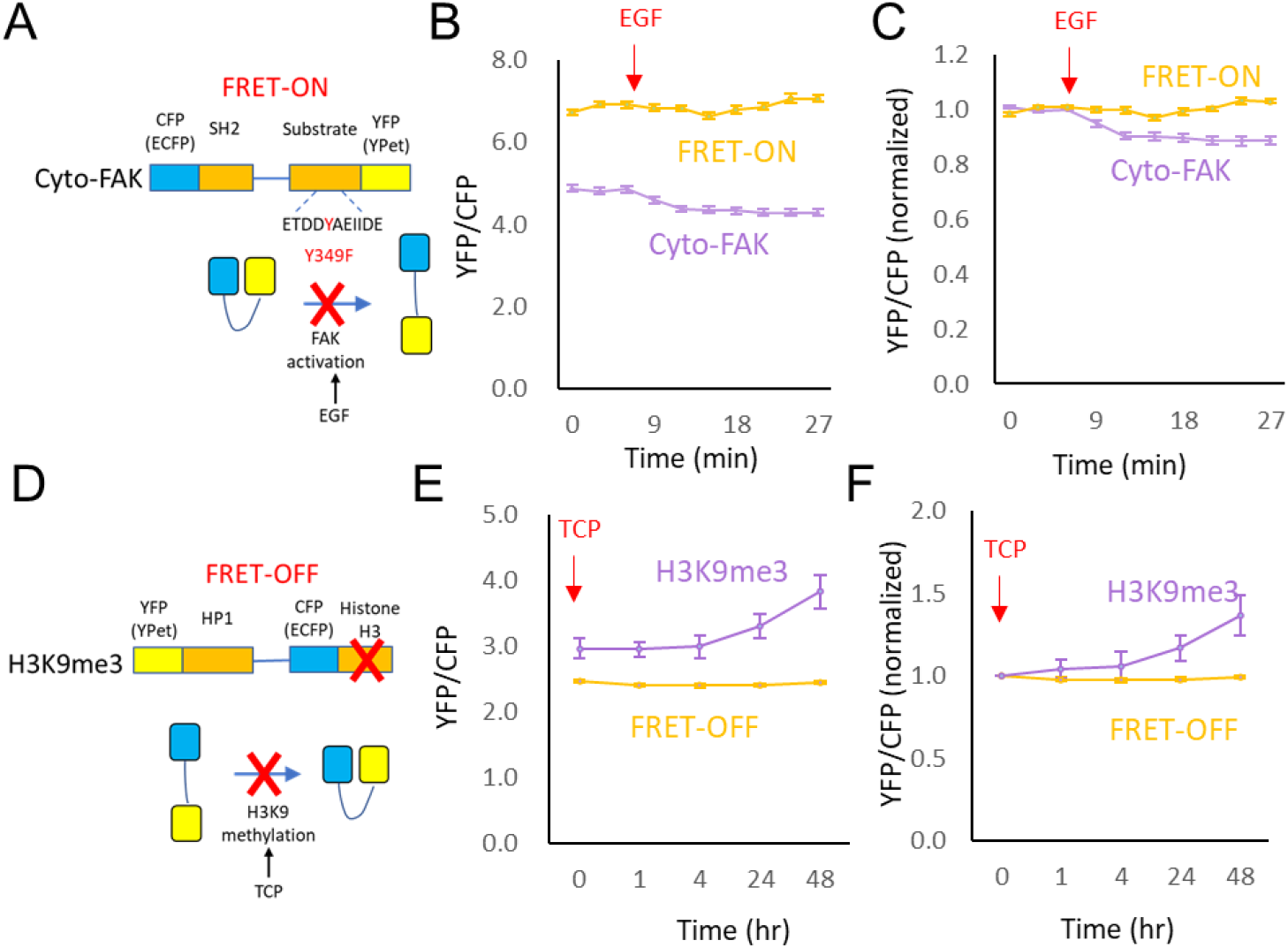
Construction of FRET calibration standards. (A) Construction of FRET-ON calibration standard by Y349F mutation of Cyto-FAK. (B) Plot of YFP/CFP ratio of CytoFAK and FRET-ON (mean ± SEM, n= 20 cells) in response to stimulation with 100 ng/ml EGF at 6 min. (C) The response in (B) normalized to the average values of the first 3 time points. (D) Construction of FRET-OFF calibration standard by removal of the H3 domain of the H3K9me9 biosensor. (E) Plot of YFP/CFP ratio of FRET-OFF and H3K9me3 (mean ± SEM, n= 20 cells) in response to 5 μM TCP over 48 hours. (F) The response in (E) normalized to the average values of the first 3 time points.

The FRET-OFF construct is based on H3K9me3 (W45A), a non-responsive control for the histone H3 lysine-9 trimethylation biosensor [23]. H3K9me3 (W45A) concontains the same donor-acceptor pair (ECFP and YPet) as the Cyto-FAK based FRET-ON. To generate the FRET-OFF construct, we removed the histone H3 domain to eliminate nuclear localization (**Fig. 2D**). While the H3K9me3 biosensor responded to the demethylase inhibitor tranylcypromine (TCP) with an increase in the YFP/CFP ratio over 48 hours, FRET-OFF showed a lower basal YFP/CFP ratio and did not respond to TCP (**Fig. 2E, F**). Together, these results suggest that FRET-ON and FRET-OFF remain in FRET and non-FRET conformations, respectively, under perturbations.

### Relative FRET ratio of FRET-ON and FRET-OFF under different excitation intensity

To validate our theoretical prediction of the FRET ratio’s dependence on excitation intensity for high-FRET species (**Fig. 1C**), we examined the relative YFP/CFP ratio of FRET-ON and FRET-OFF under various excitation intensities by adjusting the laser power. A sufficiently high excitation is expected to deplete CFP in the ground state, resulting in a non-linear relationship between fluorescence and excitation. To ensure the excitation remained within the linear range, we measured the fluorescence signal in cells expressing only CFP while varying the laser power. In our imaging setup, as the laser power for CFP excitation increased from 1% to 100%, the CFP fluorescence signal exhibited a linear increase (R^2^= 0.999) (**Fig. S2A-C**), indicating that the laser power is within the linear range for CFP excitation.

We transfected cells with either FRET-ON or FRET-OFF plasmids, each paired with barcoding proteins that are spectrally separable from CFP and YFP [17]. We mixed the FRET-ON and FRET-OFF cells for simultaneous imaging of CFP and YFP under various intensities of CFP excitation (**Fig. 3A, B**). The identity of the cells (FRET-ON vs. FRET-OFF) can be determined by the barcodes. We then calculated the relative YFP/CFP ratio between FRET-ON and FRET-OFF cells. Consistent with our theoretical modeling result (**Fig. 1C**), we found that the YFP/CFP ratio of FRET-ON normalized to that of FRET-OFF decreased monotonically with increasing excitation intensity (**Fig. 3C,D**).

**Figure 3.**
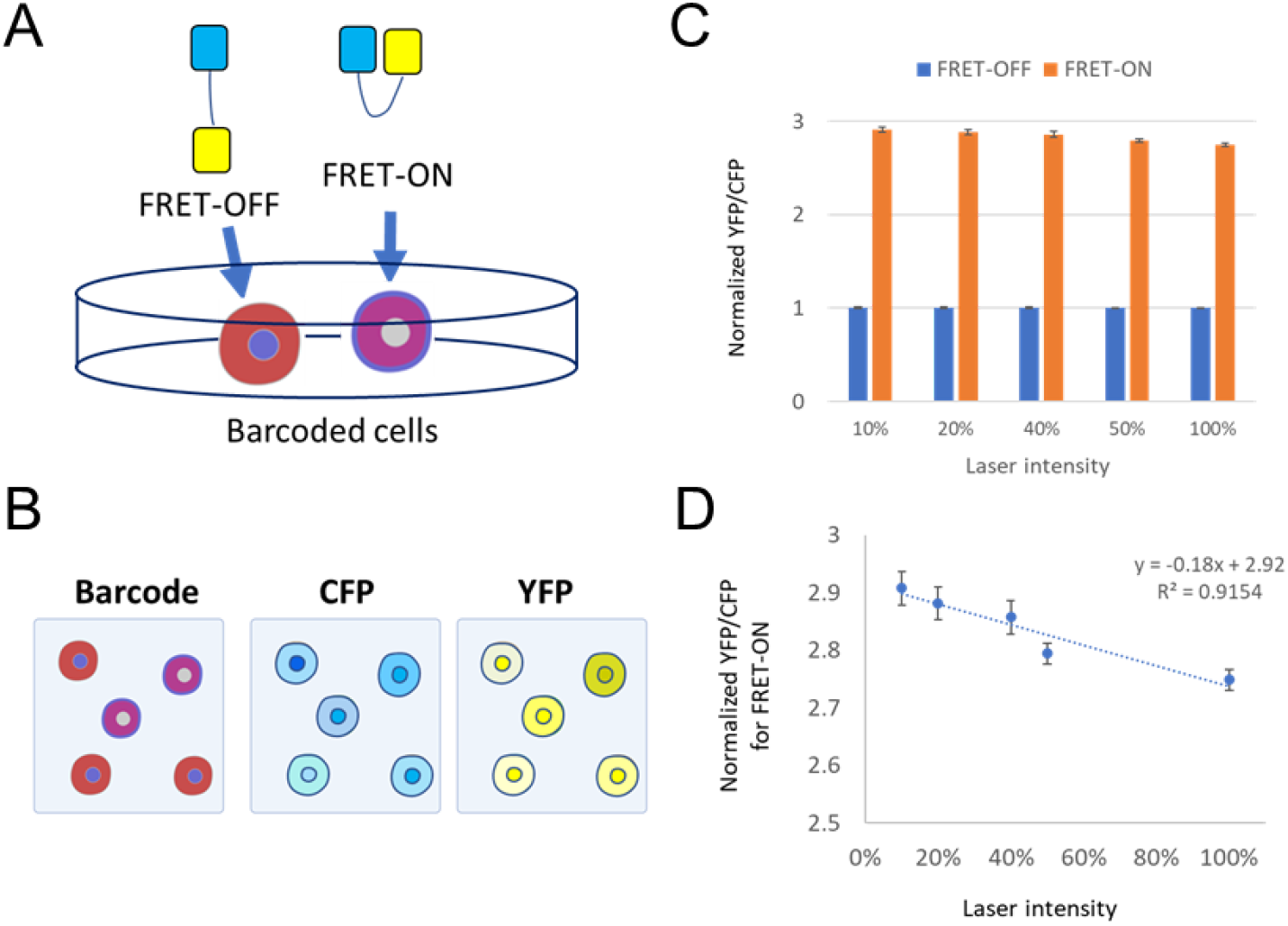
Comparison of FRET ratios of FRET-ON and FRET-OFF under different excitation intensity. (A-B) Experimental design overview. FRET-OFF and FRET-ON plasmids are introduced into cells expressing two different barcodes, which are barcoding proteins made of BFP or a red FP targeted to different subcellular locations (A). Cells are mixed and imaged for: 1) the barcodes; and 2) CFP and YFP under CFP excitation (B). (C) The YFP/CFP ratio of FRET-OFF and FRET-ON normalized to the average ratio of FRET-OFF from the same imaging experiment (mean ± SEM, n=50 cells) under various laser intensity settings. (D) Linear regression of normalized YFP/CFP for FRET-ON vs. laser intensity.

### Calibration of FRET biosensors

We next tested calibration of FRET biosensor signals using FRET-ON and FRET-OFF under different imaging settings. We chose two FRET biosensors, EKAR and Cyto-FAK, which detect the activities of ERK and FAK, respectively [22,24]. Barcoded cells expressing the four CFP-YFP constructs (FRET-ON, FRET-OFF, Cyto-FAK, and EKAR, **Fig. 4A**) were mixed and imaged under 6 settings that varied in laser power and detector gains (**Fig. 4B**). As expected, the fluorescence intensity of CFP and YFP varied widely across these different imaging conditions (**Fig. 4C**). Under the same imaging setting, YFP/CFP ratios remained relatively unaffected by the expression level (indicated by YFP fluorescence) for individual CFP-YFP constructs (**Fig. 4D,E**), except for cells with near-saturating YFP signals, which had a reduced YFP/CFP ratio. However, the YFP/CFP ratios between different imaging settings showed significant variation (**Fig. 4F**).

**Figure 4.**
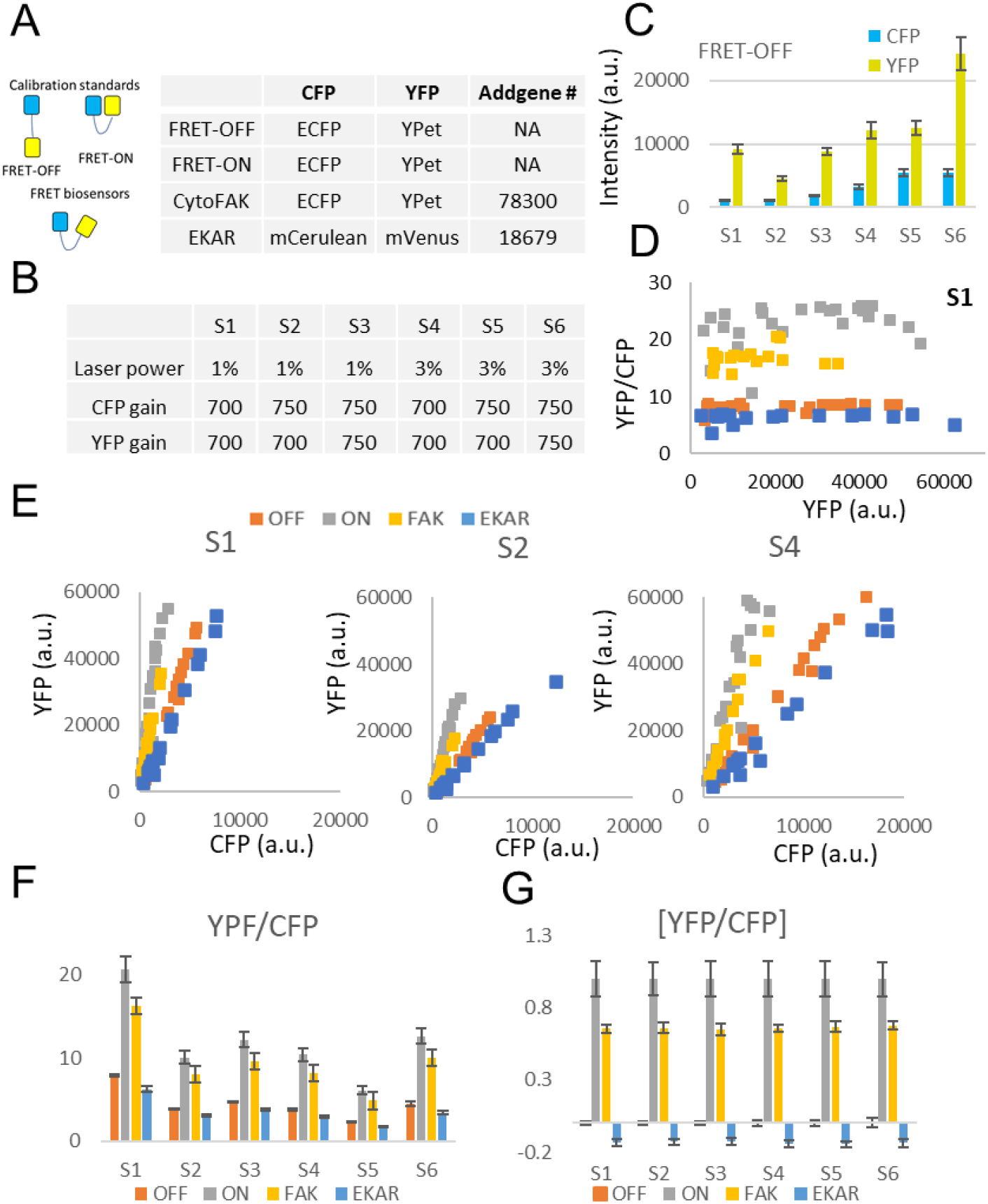
Calibration of FRET biosensors. (A) Schematic of FRET calibration standards (FRET-OFF, FRET-ON) and biosensors (CytoFAK, EKAR) as well as the CFP-YFP pair used in each construct. (B) The level of laser power and gain settings for CFP and YFP. (C) CFP and YFP levels (mean ± SEM, n= 26 cells) of FRET-OFF under different imaging settings. (D) YFP/CFP ratio of individual cells expressing the FRET constructs plotted against the YFP signal.(E) CFP vs. YFP from individual cells expressing the FRET constructs imaged at settings S1, S2, and S4. (F) YFP/CFP (mean ± SEM) across cells expressing the FRET constructs under different imaging settings. (G) YFP/CFP normalized to the calibration standards (mean ± SEM) of the FRET constructs, with FRET-OFF and FRET-ON set to 0 and 1, respectively, under different imaging settings. The cell numbers for FRET-OFF, FRET-ON, CytoFAK, and EKAR are 25, 26, 17, and 18, respectively.

Following Eq. (5), we calibrated the YPF/CFP ratios by applying a linear scaling that sets FRET-OFF and FRET-ON to 0 and 1, respectively, using the following formula:

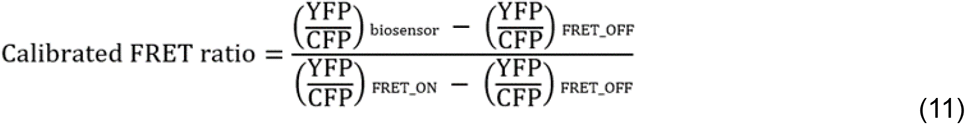

Notably, after calibration, the YFP/CFP ratios for CytoFAK and EKAR were consistent across different imaging settings (**Fig. 4G**). These findings indicate that FRET-ON and FRET-OFF can be effectively used to calibrate FRET biosensor signals under varying imaging conditions.

### Estimation of FRET efficiency

The FRET efficiency of various CFP-YFP constructs, each comprising a single copy of CFP and YFP, can be estimated by comparing the emission signals of CFP and YFP under identical imaging conditions.. As an example, we imaged cells expressing YFP, CFP, FRET-OFF, and CytoFAK under 458 nm (CFP) and 514 nm (YFP) excitation (**Fig. 5A**). (Note that although FRET-OFF and CytoFAK are used here, any two CFP-YFP fusion constructs with different FRET efficiencies can be used for the calculation.) We designated emissions at 458-500 nm (cyan) and 507-544 nm (yellow) under 458 nm emission as channels 1 and 2, respectively, and emissions at 458-500 nm (cyan) and 507-544 nm (yellow) under 514 nm emission as channels 3 and 4, respectively (**Fig. 5B**). The FRET ratio is defined as the signal ratio of channel 2 to channel 1. Additionally, we define *Y*_24_ as the signal ratio of channel 2 to channel 4 for YFP alone, and *C*_21_ as the signal ratio of channel 2 to channel 1 for CFP alone. Under our imaging settings, we found that *Y*_24_ = 0.35 and *C*_21_ = 0.39 (**Fig. 5B**). Using these numbers, we can estimate the amount of CFP loss due to FRET, as follows:

**Figure 5.**
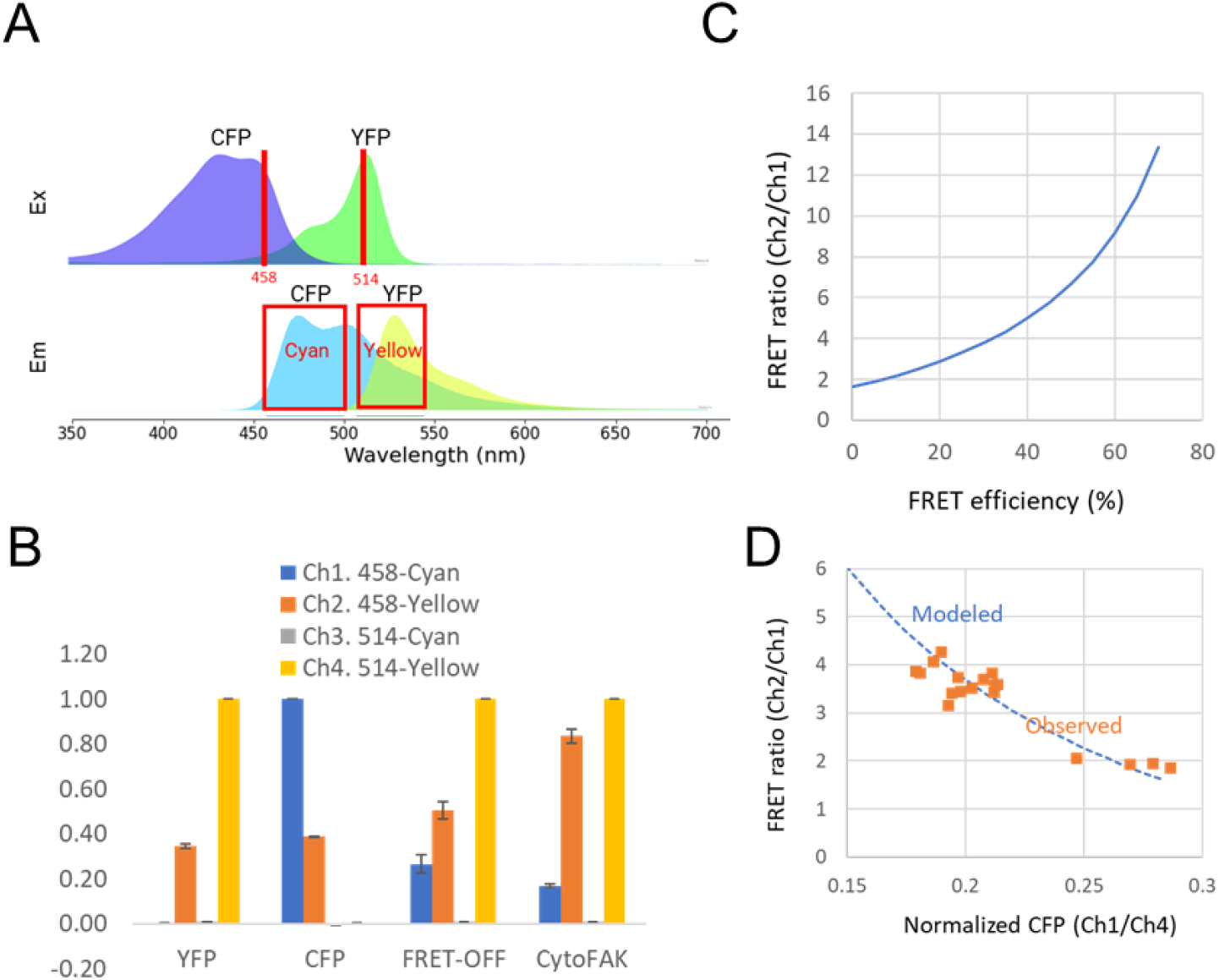
Estimation of FRET efficiency. (A) Excitation and emission spectra of CFP and YFP. The wavelengths of lasers are indicated by the red vertical lines, and the range of emission acquisition are indicated by red boxes. (B) Signals of the four channels normalized to that of the highest signal for cells expressing the four constructs (mean ± S.D., n=20 cells). (C) Plot of FRET ratio vs. FRET efficiency according to Eq. (12). (D) Predicted and observed FRET ratio vs. normalized CFP emission using the YFP level.

FRET causes a loss in CFP and an increase in YFP fluorescence. Suppose that, for a CFP-YFP construct in the absence of FRET, the CFP signal in channel 1 is *c* when the YFP signal in channel 4 is 1.00. The signal in channel 2 is *C*_21_ · *c + Y*_24_ · 1. 00. If FRET causes CFP to decrease by an amount *x* in channel 1 (i.e. the signal in channel 1 becomes *c* − *x*). In this condition, the YFP fluorescence will increase by an amount proportional to *x*. The total signal in channel 2 can be expressed as *C*_21_ · (*c* − *x*) *+ Y*_24_ · (1 *+ ax*), where *a* is a constant of proportionality.

Using the measurements in **Fig. 5B**, for FRET-OFF,

> Channel 1: *c* − *x*_*OFF*_ = 0. 27
>
> Channel 2: *C*_21_ · (*c* − *x*_*OFF*_) *+ Y*_24_ · (1 *+ ax*_*OFF*_) = 0. 50
>
> Where *x*_*OFF*_ is the decrease of channel 1 signal due to FRET

Similarly, for CytoFAK,

> Channel 1: *c* − *x*_*FAK*_= 0. 17
>
> Channel 2: *C*_21_ · (*c* − *x*_*FAK*_) *+ Y*_24_ · (1 *+ ax*_*FAK*_) = 0. 84
>
> Where *x*_*FAK*_is the decrease of channel 1 signal due to FRET

Solving the system of equations gave:

> *c* = 0. 282
>
> *x*_*OFF*_ = 0. 012
>
> *x*_*FAK*_= 0. 112
>
> *a* = 10. 81

In other words, FRET-OFF had a loss of CFP of 0.012/0.282 = 4.2%, whereas CytoFAK had a loss of CFP of 0.112/.282 = 39.7%

The relationship between CFP loss and the FRET ratio (channel 2/channel 1) is:

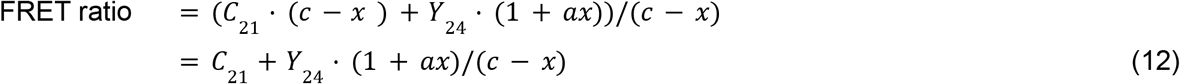

Note that FRET efficiency, represented by *x*/*c*, is the fraction of energy transferred from the FRET donor (CFP) to the acceptor (YFP). Using the data from this experiment, we plotted the FRET ratio against FRET efficiency (**Fig. 5C**). The FRET ratio becomes increasingly sensitive to changes in FRET at higher levels, approaching infinity as FRET efficiency nears 100%. Cells with higher FRET ratios are also expected to have reduced CFP fluorescence. This can be seen by plotting the FRET ratio (channel 2/channel 1) vs. normalized CFP emission using YFP level (channel 1/channel 4). The predicted and observed FRET ratio vs. normalized CFP emission is shown in **Fig. 5D**, in which the observed values are derived from mixed populations of cells expressing FRET-OFF or CytoFAK. Together, these results demonstrate that simultaneous imaging of cells expressing CFP-YFP constructs along with CFP or YFP can effectively determine FRET efficiency.

## Discussion

While FRET biosensors are versatile and powerful tools for studying biological processes in real time, their ability to provide a comprehensive view of molecular networks over extended time periods is limited by challenges in multiplexing and complex calibration processes. As a result, they are typically used for monitoring individual processes over relatively short durations. In this study, we addressed these limitations by introducing calibration standards in a subset of cells, using a recently developed biosensor barcoding method for multiplexed imaging of biosensors.

Our theoretical analysis revealed that the FRET ratio (the ratio of acceptor to donor signal) is dependent on excitation intensity. Therefore, calibrating the FRET ratio across different excitation intensities requires standards in both low and high FRET states. We validated this prediction by constructing FRET-OFF and FRET-ON calibration standards, and showed that calibrated FRET ratios of biosensors remain consistent regardless of imaging conditions.

Together, our results suggest a strategy for calibrated, highly multiplexed imaging of FRET biosensors using biosensor barcoding (**Fig. 6**). In this strategy, FRET-ON and FRET-OFF calibration standards, along with various biosensors, are separately introduced into cells expressing different barcoding proteins, which are spectrally distinct FPs targeted to specific subcellular locations (**Fig. 6A**). These different cell populations are then combined for simultaneous imaging of the barcodes, along with CFP and YFP fluorescence under CFP excitation. Cells expressing biosensors and the calibration standards are identified through their associated barcodes (**Fig. 6B**). The YFP/CFP ratios for the standards and biosensors are calculated, and the values of the standards are used to calibrate those of the biosensors (**Fig. 6C**). This calibration allows for direct comparison of the YFP/CFP ratios for each biosensor across different timepoints or experiments. Moreover, if YFP and CFP are included in the mix, the actual FRET efficiency can be calculated.

**Figure 6.**
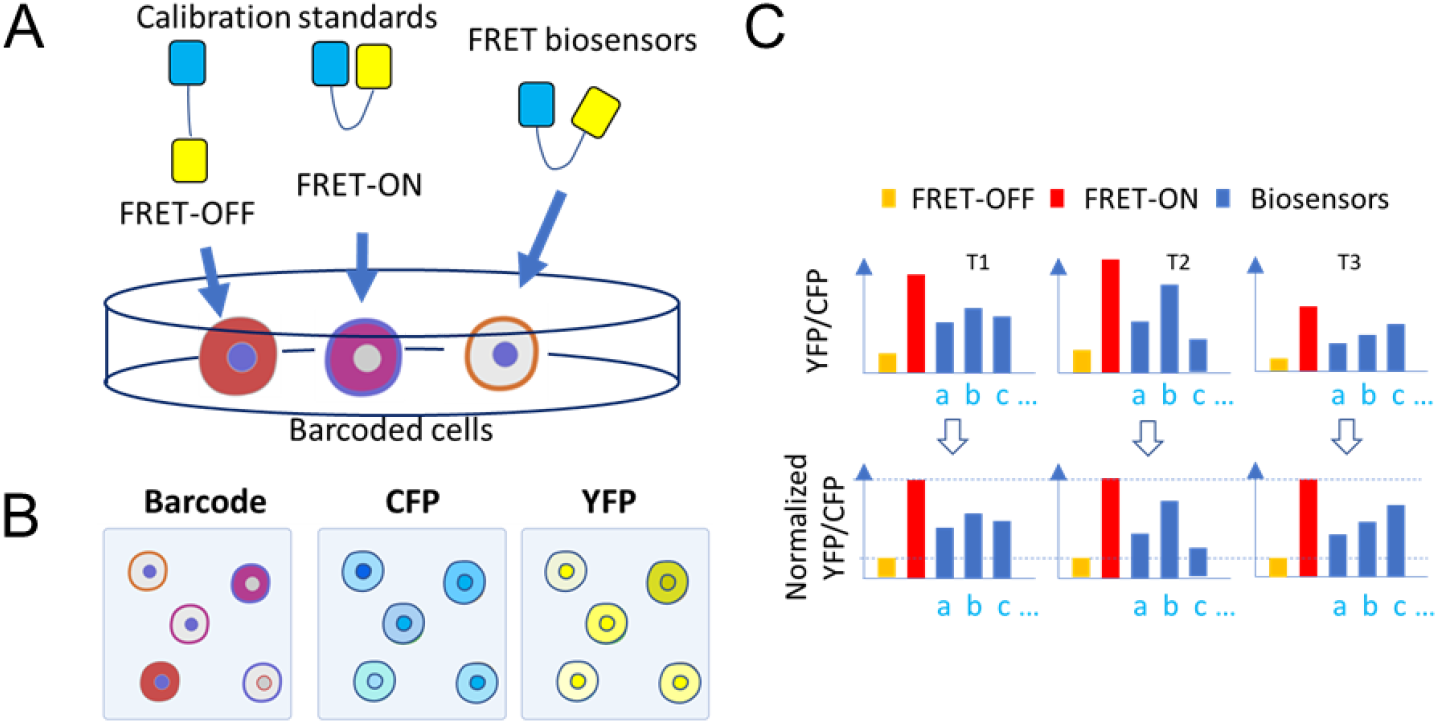
Strategy for calibrated and highly multiplexed imaging of FRET biosensors. (A) Calibration standards (FRET-OFF and FRET-ON) and biosensor plasmids are introduced into cells expressing different barcodes. (B) Cells are mixed and imaged for: 1) the barcodes; and 2) CFP and YFP under CFP excitation. (C) The YFP/CFP ratio of calibration standards and biosensors are calculated for images taken at different time points (T1-T3). The YFP/CFP of FRET-ON and FRET-OFF calibration standards are used to normalize those of the biosensors using Eq. (12).

The principle presented in this study can be broadly applied to the calibration of other ratiometric biosensors. For example, biosensors such as the ATP sensor iATPsnFR or the NADH/NAD sensor Peredox [25,26], are designed by linking a cpGFP, which responds to the target ligand, with a red FP (e.g. mCherry). The intensity of a cpGFP is normalized to that of the red FP as the readout for target activities. Since these biosensors do not depend on FRET, a single calibration standard made of a GFP-RFP fusion protein is sufficient for calibrating the relative fluorescence of the two FPs under different imaging conditions.

In conclusion, our method provides a straightforward approach for calibrated and multiplexed measurement of biosensors across diverse applications, enabling consistent comparisons between experiments and supporting long-term imaging studies.

## Materials and Methods

### Cell culture

HeLa and HEK293T cells, purchased from ATCC, were grown at 37°C, 5% CO_2_ in DMEM high glucose medium (Gibco, #11965092) supplemented with 10% FBS (Corning Cellgro, 35-010-CV), 1 mM sodium pyruvate (Gibco, #11360070), and 1X nonessential amino acids (Gibco, #11140076).

### Chemical reagents

The EGF stock solution was prepared by dissolving EGF (Sigma-Aldrich, E9644) in 10 mM acetic acid to a final concentration of 1 mg/ml. TCP stock solution was prepared by dissolving TCP in DMSO to a final concentration of 5 mM. All stocks were stored at -20°C.

### Plasmid construction

Plasmids for transient expression were obtained from Addgene: EKAR (#18679), CytoFAK (#78300), H3K9me3 biosensor (#120802), and H3K9me3 (W45A) biosensor (#120808). For lentiviral vector versions of these biosensors, the Gateway cloning system was used to first generate the entry vectors and then the destination vectors. Due to the highly similar DNA sequences of the two fluorophores in EKAR (mCerulean and mVenus), we replaced them with the fluorophore pair from CytoFAK (ECFP and YPet) and synthesized the modified gene, named EKAR-1 (GenScript). To create entry vectors, we designed a pair of PCR primers with an attB1 site at the 5’ end and an attB2 site at the 3’ end to amplify the biosensor genes using EKAR-1, CytoFAK, the H3K9me3 biosensor, and the H3K9me3 (W45A) biosensor as templates. The PCR products were cloned into the pDONR221 vector (Invitrogen, #12536017) using Gateway BP Clonase II (Invitrogen, #11789020). The resulting entry vectors were then recombined with the pLenti-CMV-Neo destination vector (Addgene, #17392) to produce lentiviral expression vectors using Gateway LR Clonase II (Invitrogen, #11791020). The resulting plasmids can be used either for transient transfection or as transfer plasmids for lentivirus production.

### Cell transfection

Cells were transiently transfected using the GenJet™ In Vitro DNA Transfection Reagent (Ver. II) following the manufacturer’s protocol (SignaGen Laboratories, #SL100489). Briefly, 4×10^5^ cells were seeded in one 3.5 cm culture dish the day before transfection. On the day of transfection, the culture medium was replaced with 1 mL fresh complete medium without antibiotics. The GenJet-DNA complex was prepared by diluting (1) 1 μg of DNA in 50 μL serum-free DMEM, and (2) 3 μL GenJet reagent in 50 μL serum-free DMEM in two separate tubes, and then mix the two together by adding GenJet into DNA. The complex was allowed to form at room temperature for 15 min, and then added dropwise onto the cells. Target protein expression in cells can be observed 24 to 48 h post transfection.

### Microscopy

Cells were transfected with plasmids encoding the barcodes and CFP-YFP constructs (i.e. calibration standards or biosensors) in separate wells. Prior to imaging experiments, cells transfected with different barcode/CFP-YFP constructs combinations were harvested, mixed, and reseeded. Microscopy was carried out on either a Zeiss 780 or 880 confocal microscope equipped with a spectral detector (for imaging barcoding proteins) and a motorized stage (for capturing multiple viewing fields). The detailed imaging protocol has been described in our earlier publication [18]. Briefly, for the barcodes, BFP images were acquired under 405 nm excitation and spectral images (550-700 nm) of the red FPs under 561 or 633 nm excitation. For the CFP-YFP constructs, CFP and YFP images were acquired under 457 nm excitation.

### Image analysis

Images of barcodes and CFP-YFP constructs were processed and analyzed with NIH ImageJ and Fiji [27,28]. Cells were manually segmented to identify individual cells. The barcode for each cell was determined based on the color and subcellular localization of the barcoding proteins, allowing for the identification of the specific CFP-YFP constructs expressed in each cell. Pixel values for YFP and CFP were measured, and the YFP/CFP ratios were calculated using Microsoft Excel. Microsoft Excel was also used for statistical analysis and to generate graphs.

## Author Contribution

J.M.Y. and C.H.H. conceived the project. G.W.C. carried out theoretical modeling. J.W.W., J.M.Y., C.C.C., G.A., and S.W. conducted the experiment. J.W.W., J.M.Y., C.C.C. and C.H.H. analyzed the data. J.M.Y., G.W.C., and C.H.H. wrote the manuscript with input from all authors. J.M.Y. and C.H.H. supervised the project.

## Acknowledgments

This work was supported by NIH grants R01GM136711 (to C.H.H.), Cervical Cancer SPORE P50CA098252 Career Development Award (to J.M.Y.) and Pilot Project Award (to C.H.H.), and the Sol Goldman Pancreatic Cancer Research Center (to C.H.H.).

