## Supplemental Figures for "Calibration of FRET-based biosensors using multiplexed biosensor barcoding"

Wu et al.

#### Figure S1

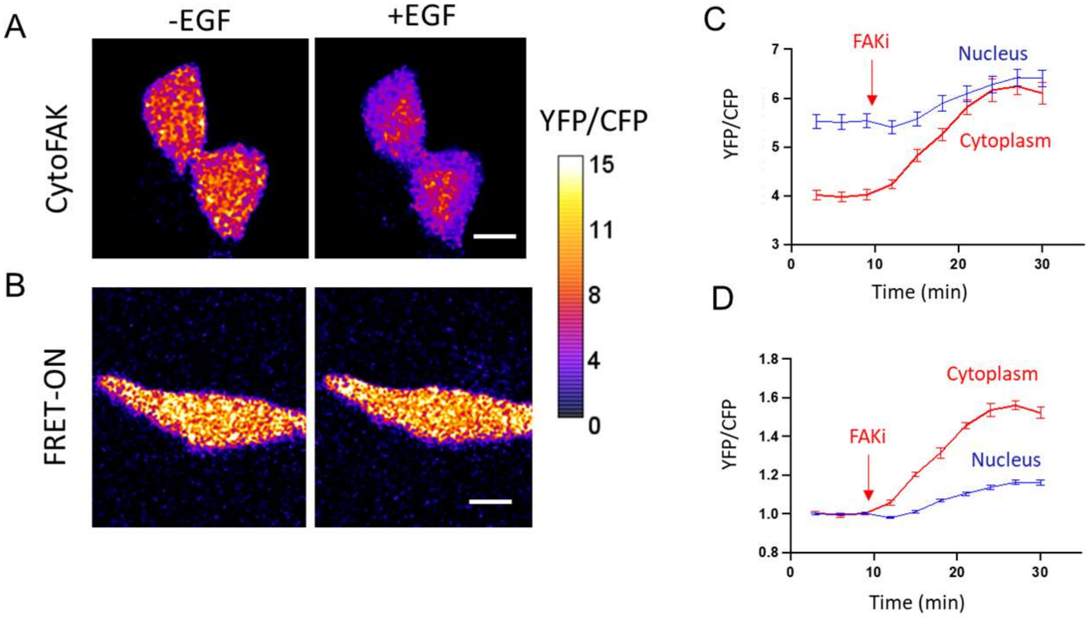


**Figure S1. Responses of CytoFAK and FRET-ON**

(A-B) YFP/CFP ratio images of CytoFAK (A) and FRET-ON (B) before and after treatment with 100 ng/ml EGF. Scale bar: 10 μm.

(C) Plot of YFP/CFP ratio of the nuclear and cytosolic regions in cells expressing CytoFAK (mean ± SEM, n= 20 cells each) in response to the FAK inhibitor PF-562271. (D) The response in (C) normalized to the average values of the first 3 time points.

### Figure S2

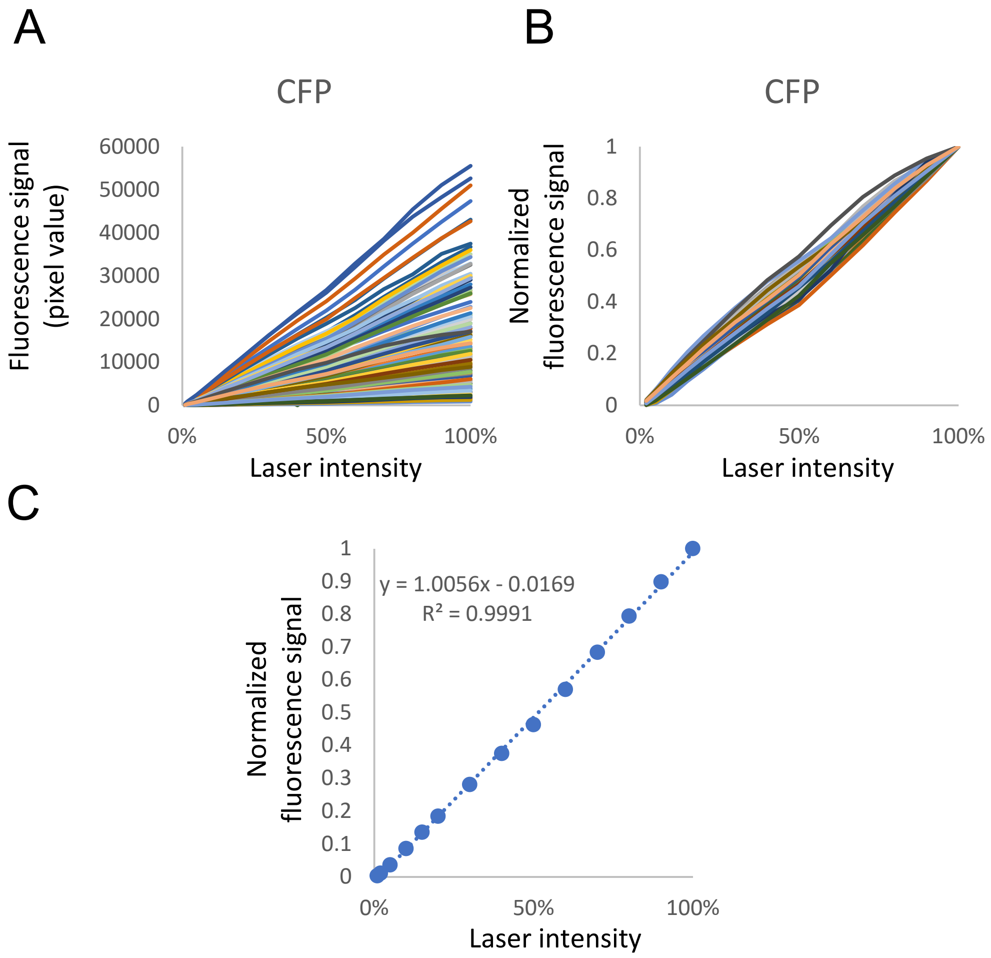


**Figure S2. Linear relationship between CFP fluorescence and laser intensity.**
(A) Fluorescence signals from 200 cells expressing CFP under laser intensity between 1-100%. (B) Fluorescence signals of the cells in (A) normalized to the intensity at 100% laser. (C) Quantification of the correlation between the mean normalized fluorescence signal vs. laser intensity.
